# HiC-ACT: Improved Detection of Chromatin Interactions from Hi-C Data via Aggregated Cauchy Test

**DOI:** 10.1101/2020.10.28.359869

**Authors:** Taylor M. Lagler, Yuchen Yang, Armen Abnousi, Ming Hu, Yun Li

## Abstract

Genome-wide chromatin conformation capture technologies such as Hi-C are commonly employed to study chromatin spatial organization. In particular, to identify statistically significant long-range chromatin interactions from Hi-C data, most existing methods such as Fit-Hi-C/FitHiC2 and HiCCUPS assume that all chromatin interactions are statistically independent. Such an independence assumption is reasonable at low resolution (e.g., 40Kb bin), but is invalid at high resolution (e.g., 5 or 10Kb bins) since spatial dependency of neighboring chromatin interactions is non-negligible at high resolution. Our previous hidden Markov random field based methods accommodate spatial dependency but are computationally intensive. It is urgent to develop approaches that can model spatial dependence, in a computationally efficient and scalable manner. Here, we develop HiC-ACT, an aggregated Cauchy test (ACT) based approach, to improve the detection of chromatin interactions by post-processing results from methods assuming independence. To benchmark the performance of HiC-ACT, we re-analyzed deeply sequenced Hi-C data from a human lymphoblastoid cell line GM12878 and mouse embryonic stem cell line (mESC). Our results demonstrate advantages of HiC-ACT in improving sensitivity with controlled type-I error. By leveraging information from neighboring chromatin interactions, HiC-ACT enhances the power to detect interactions with lower signal to noise ratio and similar (if not stronger) epigenetic signatures that suggest regulatory roles. We further demonstrate that HiC-ACT peaks show higher overlap with known enhancers than Fit-Hi-C/FitHiC2 peaks, in both GM12878 and mESC. HiC-ACT, effectively a summary statistic based approach, is computationally efficient (~6 minutes and ~2GB memory to process 25,000 pairwise interactions).

## Introduction

Chromatin spatial organization plays a critical role in genome functions such as transcription regulation and DNA replication^1–3^. Studies have shown that millions of putative *cis*-regulatory elements, such as enhancers, exist within the genome, many of which are far away in one-dimensional (1D) genomic distance from their target genes (e.g. up to Mb away)^1,2,4–6^. Due to the abundance of enhancers and their long-range regulation, systematic mapping of enhancer-promoter interactions is challenging^1^.

Genome-wide chromosome conformation capture techniques such as Hi-C^7^ have been widely used to study three-dimensional (3D) organization of chromatin. Hi-C data can be summarized into contact matrix of all possible pairwise interactions between ligated fragments genome-wide. As comprehensive chromatin interaction maps become increasingly prominent due to increases in sequencing capacity and decreases in cost, there is an urgent need to develop tools to analyze and interpret this data^8^. Such methods to detect statistically significant long-range chromatin interactions (also referred to as “peak callers”) seek to determine if the observed contact frequency is significantly higher than expected from chromatin random collision.

Fit-Hi-C^9^ is a popular method to evaluate pairs of chromatin loci independently, and assigns each pair a statistical confidence (*p*-value). Fit-Hi-C corrects distance dependence and potential systematic biases in Hi-C datasets by fitting non-parametric splines to model the background chromatin contact frequency^9–11^. Recently, a re-implementation, FitHiC2^10^, was released. Along with the addition of new computational modules, FitHiC2 can be applied to the highest-resolution Hi-C datasets currently available^10^. However, in high-resolution data (e.g. 5Kb or 10Kb bin resolution), neighboring chromatin interactions are unlikely to be independent, as assumed in FitHiC2. When this independence assumption is violated, the *p*-values corresponding to chromatin interactions is inaccurate.

We have previously demonstrated that spatial dependency is non-negligible when analyzing Hi-C data at high resolution. Accordingly, we developed hidden Markov random field (HMRF) based methods HMRF-Bayes^12^ and FastHiC^13^ to accommodate spatial dependency for improved statistical properties. However, compared with FitHiC2 that analyzes each pair of chromatin segments separately, our HMRF based framework is more computationally intensive.

HiCCUPS^14^ (Hi-C Computational Unbiased Peak Search) is another commonly adopted method for identifying significant chromatin interactions. Unlike Fit-Hi-C/FitHiC2 and our HMRF based methods which use a global background model^9,10,12^, HiCCUPS uses a local background model where each chromatin loci pair has a unique model influenced by information from local neighborhoods^14^. This model defines peaks based on whether the loci pair interacts significantly more than loci pairs in its neighborhood. Therefore, HiCCUPS effectively detects summits of chromatin interactions, rather than peaks as in Fit-Hi-C/FitHiC2 and our HMRF based methods. A most recently published method, MUSTACHE, similarly relies on a local background model and detects summits using a scale-space modeling framework enlightened by methods in computer vision^15^. The summit-detection strategy is valuable in distinguishing the most frequently interacting pairs from its neighborhood, but limits its ability to identify many functionally important interactions linking regulatory elements such as promoters and enhancers^10,14^.

In this paper, we develop HiC-ACT, a method for post-processing peak calling results from methods that do not consider spatial dependency. HiC-ACT’s post-processing via an aggregated Cauchy test approach accounts for possible correlation between adjacent loci pairs from high resolution Hi-C data. HiC-ACT is flexible in application, only requiring the input of bin identifiers and corresponding raw *p*-value, and allows users to specify a smoothing parameter based on the data resolution. Moreover, HiC-ACT does not require any information about the underlying correlation structure in the data while being able to account for the inherent correlation between bin (loci) pairs.

The implementation of p-value smoothing in HiC-ACT improves identification of significant chromatin interactions and recovers information lost in sparse data. Since HiC-ACT borrows information from neighboring loci pairs, it calls peaks rather than summits. Thus, we chose to compare HiC-ACT to FitHiC2. Both simulation studies and real data analysis demonstrate that HiC-ACT outperforms FitHiC2 in increasing recall with comparable precision.

In the remainder of this article, we will specify the HiC-ACT model and provide details regarding the workflow. Next, we show real data-based simulation results based on Hi-C data from the human lymphoblastoid cell line GM12878 at various sequencing depths. Then, we perform real data analysis using Hi-C datasets from GM12878 and mouse embryonic stem cells (mESC). Finally, we conclude with some discussions.

## Material and Methods

### Aggregated Cauchy Combination Test

HiC-ACT is based on the aggregated Cauchy combination test^16^ to combine a set of *p*-values *p*_1_, *p*_2_,…, *p_k_*. We use a linear combination of transformed *p*-values with non-negative weights:

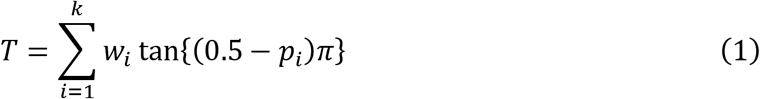

where *p_i_* is the individual *p*-value and *w_i_* is the non-negative weight such that 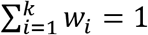.

When only one *p*-value is considered (*k* = 1), it is straightforward to show that *T* follows a Cauchy distribution (location parameter *x*_0_ = 0, scale parameter *γ* = *w*_1_ = 1) under the null hypothesis that *p*_1_ is uniformly distributed between 0 and 1 (Appendix).

In an application to rare variant association analysis, Liu et al. showed that this combination of p-values, *T*, follows a standard Cauchy distribution (*x*_0_ = 0, *γ* = 1) under the null hypothesis^17^.

Assume that the *p*-values are calculated from Z-scores and let **X** = (*X*_1_, *X*_2_,…, *X_k_*)^*T*^, were *X_i_* is a test statistic corresponding to *p_i_*. The null hypothesis can then be written as *H*_0_: E[**X**] = **0**.

Thus, *p_i_* can be expressed as *p_i_* = 2(1 – *Φ*(|*X_i_*|)). We can then re-write *T* (Equation 1) as:

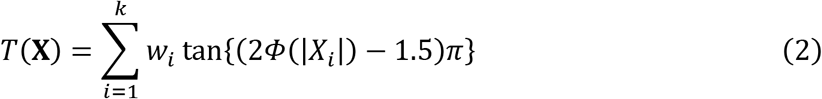

If the *p_i_*’s are perfectly dependent (i.e. all the *p_i_*’s are equal or linear functions of one another) or perfectly independent, then it can be shown easily that the sum of two independent Cauchy random variables is also Cauchy. Furthermore, it has been shown that this holds even when the *p_i_*’s are correlated^16,17^.

Liu et al. further showed that under arbitrary dependency structures *T*(**X**) has approximately a Cauchy tail^16,17^. They also demonstrated that when the p¿’s are correlated, it has very limited effect on the tail of the distribution. Consequently, we can transform the test statistic *T* back to a *p*-value using Cauchy(0,1).

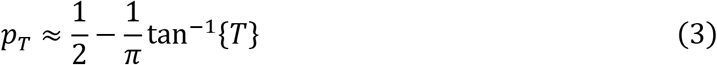

This approximation is particularly appealing for small *p*-values, which are typically observed in Hi-C data analysis. Liu also argued that if the individual *p*-values are conservative, *p_T_* will be conservative as well and that the Type I error is controlled.

### HiC-ACT Test Statistic

Using the framework above, we specify the HiC-ACT statistic as follows. Let *p_ij_*-represent the *p*-value for chromatin interaction between bin *i* and bin *j* from a specific Hi-C peak calling method. Consider the null hypothesis that the contact frequency between bin pair (*i, j*) is due to random chromatin collision. Define the HiC-ACT test statistic *T_ACT_ij__* as

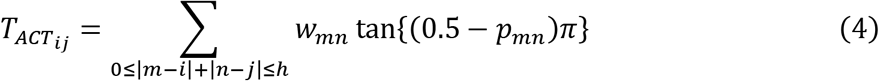

Here, *h* is the local smoothing bandwidth. We followed the strategy adopted by the HiCRep^18^ method to determine the size of the smoothing window based on data resolution (Table S1). We take *w_mn_* to be the Gaussian kernel weight function, defined as

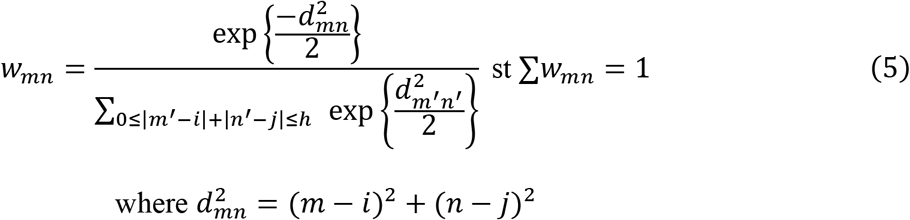

Note that the *p*-value for the bin pair itself contributes to the statistic, and thus the smoothed *p*-value.

Based on the theory established, *T_ACT_ij__* approximately follows a standard Cauchy distribution under the null hypothesis. Therefore, the *p*-value for *T_ACT_ij__* can be approximated by

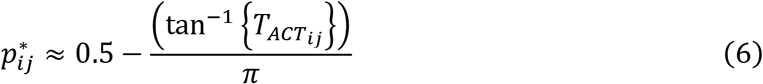

We can interpret 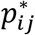 as the local neighborhood smoothed *p*-value. Intuitively, for a biologically meaningful chromatin interaction, all bin pairs in its neighborhood are more likely to have significant *p*-values. Thus, the combined *p*-value 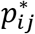 tends to be more significant and is driven by small *p*-values in its neighborhood. Through the properties of the aggregated Cauchy combination test, HiC-ACT specifically accounts for the inherent correlation between the contact frequency of neighboring pairs and maintains the benefit of no correlation structure assumption.

### Workflow

To implement HiC-ACT, we first obtain results from a standard peak caller not considering spatial dependency. HiC-ACT only requires bin pair identifiers and corresponding *p*-values. Next, we set *h* based on the data resolution (see Table S1 for suggestions). Then, we identify a set of (*i, j*) bin pairs of interest, e.g. by selecting if *p_ij_* is less than a specified threshold. We recommend that this threshold depends on the number of pairwise raw reads in the data (Table S2). For each (*i, j*) pair, HiC-ACT determines all possible (*m, n*) pairs that meet criterion in Equation 4, calculates the weights, then computes *T_ACT_ij__* and its corresponding *p*-value 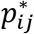 for each (*i, j*) pair in the set of interest using Equation 6.

In Figure 1, we present a motivating example for HiC-ACT using 10Kb GM12878 Hi-C data acquired from the Rao et al. study consisting of ~4.9 billion pairwise contacts^14^. Each colored pixel on the heatmap represents the strength of the FitHiC2 interaction (*p*-value), represented on a −log10 scale. This specific chromatin interaction (i.e. bin pair) is centered at (50,625,000, 50,975,000) on chromosome 22 (marked by a blue ‘x’) has one end overlapping with an identified super enhancer from the Roadmap Epigenomics Consortium^19^ and one end overlapping with the transcription start site (TSS)^20^ of the highly expressed *TRABD* gene (FPKM = 17)^21^. However, when the data is down-sampled to ~1 billion raw reads (a more realistic sequencing depth), this interaction is not classified as a peak by FitHiC2 (*p* = 8.62e-04) (see Table S2 for details on how peaks were determined) (Figure 1A). When HiC-ACT is applied to these FitHiC2 calls, the resulting *p*-value is highly significant (*p* = 2.73e-19) as expected given the biological evidence. Figure 1B displays the corresponding heatmap for FitHiC2 interactions/*p*-values called on the full GM12878 data (~4.9 billion reads). The FitHiC2 *p*-value here for the specified interaction is 3.50e-11. Comparing Figure 1A to Figure 1B, we notice that information is lost in data with shallower sequencing depth. HiC-ACT is able to recover some information lost in Hi-C data with shallower sequencing depths by leveraging information from neighboring loci.

**Figure 1.**
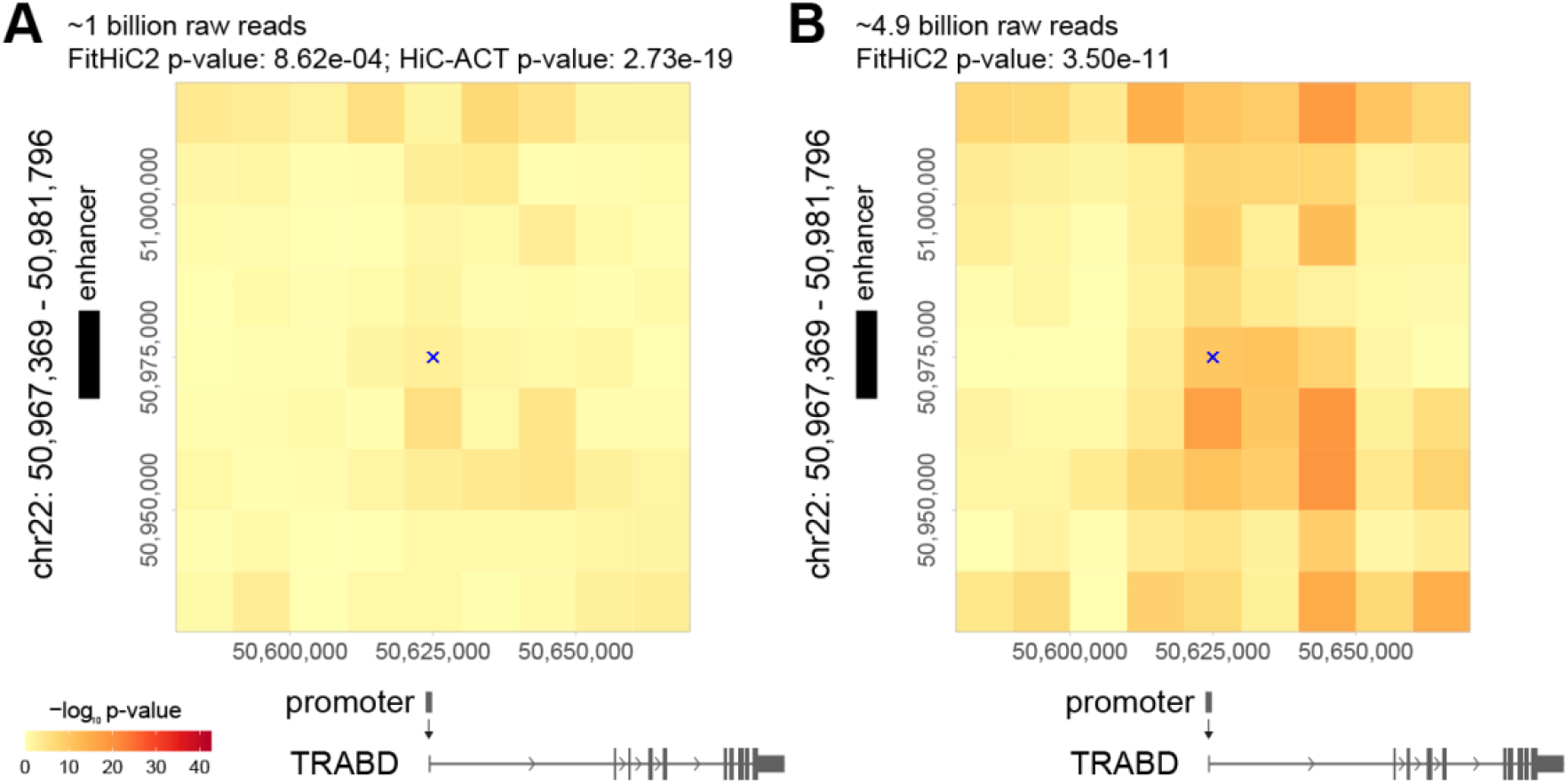
Motivating Example for HiC-ACT. FitHiC2 was applied to the GM12878 10Kb Hi-C data down-sampled to ~1 billion raw reads **(A)** as well as to the full GM12878 data (~4.9 billion raw reads) **(B)**. Each colored pixel on the heatmap represents the strength of the FitHiC2 interaction (*p*-value), represented on a −log10 scale. Here, the chromatin interaction (i.e. bin pair) of interest is centered at (50,625,000, 50,975,000) on chromosome 22 (marked by a blue ‘x’). This interaction has one end overlapping with an identified super enhancer from the Roadmap Epigenomics Consortium (marked by black bar on the left side) and the other end overlapping with the transcription start site (TSS) for the TRABD gene, indicating a possible functional interaction. However, this interaction is not marked as significant by FitHiC2 in the lower sequencing depth (*p* = 8.62e-04) **(A)**. When HiC-ACT is applied to these FitHiC2 calls, the resulting p-value is highly significant (*p* = 2.73e-19) as expected given the biological evidence. By using information from neighboring loci, HiC-ACT is able to recover some information lost in Hi-C data with shallower sequencing depths.

## Results

### Real Data-Based Simulations

We first used real data-based simulations to assess the performance of HiC-ACT. The simulations were based on the 10Kb GM12878 (human lymphoblastoid cell line) Hi-C data consisting of ~4.9 billion pairwise contacts^14^. FitHiC2 results generated from this high-depth data were treated as the truth. Approximately 1.57 million significant chromatin interactions were identified based on the criterion that the observed contact count > 15, the expected contact count > 5, the ratio of observed to expected > 1.5, and the *p*-value < 1.0e-12. The *p*-value threshold was informed by a recent study of 10Kb bin resolution deeply sequenced Hi-C data from human brain cortex, where high-confidence regulatory chromatin interactions were determined using *p* < 2.31e-11^5^.

To simulate more realistic sequencing depths, we down-sampled the GM12878 Hi-C data to 10-40% of the original data size corresponding to ~0.5-2.0 billion raw reads. For each of these down-sampled data, we ran FitHiC2 then applied HiC-ACT. Following HiCRep^18^, we chose the smoothing bandwidth (*h*) to be 20 since we analyzed the data at 10Kb resolution.

Significant pairwise interactions were defined using sequencing depth-specific threshold of minimum observed contact count, minimum expected contact count, global significant *p*-value threshold, and for HiC-ACT, initial *p*-value filtering. In each case, a minimum ratio between observed count and expected count of 1.5 was required to determine a significant interaction. Table S2 provides recommendations for defining significant interactions (i.e. peaks) based on this simulation using sequencing depth-specific global *p* threshold.

Using the significant interactions defined from the full GM12878 data as the truth, we counted the number of interactions correctly classified as significant or insignificant by HiC-ACT and FitHiC2. HiC-ACT correctly identified 80-400% more significant interactions than FitHiC2 and achieves comparable precision (Table 1, Table S3).

**Table 1.**
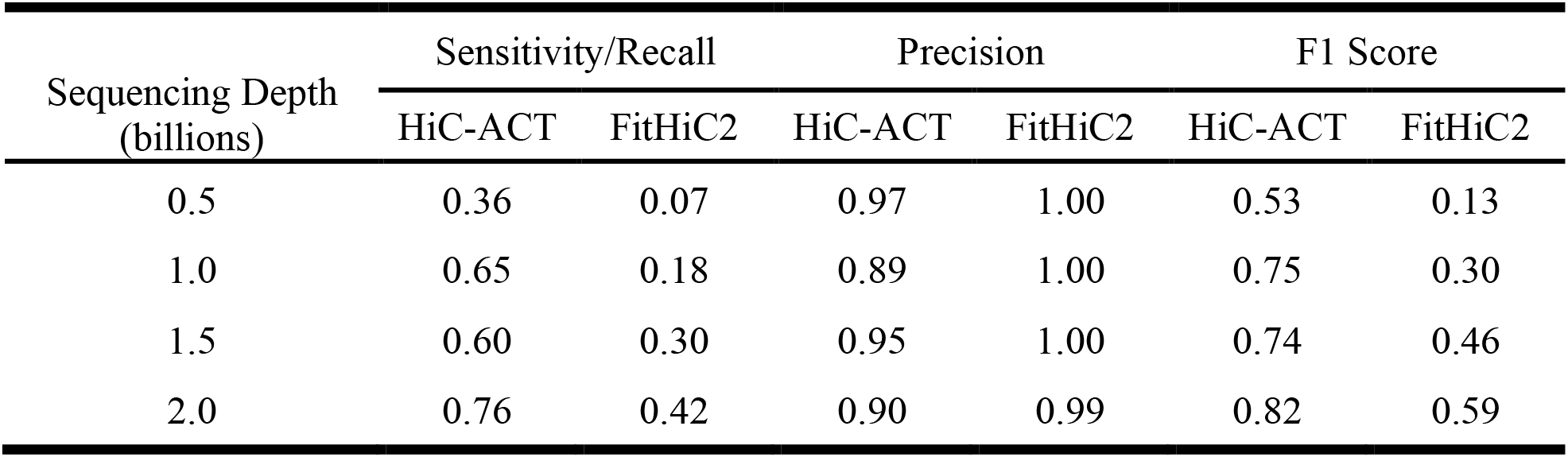
HiC-ACT considerably improves sensitivity with affordable loss of precision. Sensitivity, precision, and corresponding F1 score (harmonic mean of precision and recall) of calling true peaks at various GM12878 10Kb sequencing depths (in approximate billions of raw reads) is reported. Peaks are defined using the guidelines in Table S2 and peaks called by FitHiC2 in the full GM12878 data (~4.9 billion raw reads) are treated as truth.

While HiC-ACT tends to be driven by the most significant interactions in a neighborhood (those pairs with extremely small *p*-values), it maintains large *p*-values for truly insignificant interactions. To demonstrate this, we calculated the sensitivity/recall and precision for correct identification of significant interactions for each method. We also report the F1 score, defined as the harmonic mean of precision and recall, where a value of 1 indicates perfect precision and recall. Table 1 displays a summary of these results at various sequencing depths (in billions of raw reads). HiC-ACT considerably improves sensitivity with affordable loss of precision, as demonstrated by greater F1 scores, in all sequencing depths, though the largest improvements are seen when sequencing depth is low. We note that the pattern of these results holds when the global significance threshold is adjusted (Table S3).

In the Hi-C peak calling problem, the number of true positives (significant interactions/peaks) and true negatives (insignificant interactions/background noise) is highly unbalanced. Due to the large proportion of true negatives, comparing sensitivity versus specificity is not ideal. Precision versus recall is a more appropriate performance metric in this scenario^22^. Accordingly, specificity is omitted from Table 1 since the values for both methods are nearly 1 due to the large number of non-peaks. Specificity, along with peak classification counts can be found in Table S3.

We can further examine the relationship between true positives (i.e. correctly identifying significant interactions/calling true peaks) and false positives (i.e. incorrectly identifying interactions as significant/calling false peaks) through precision recall curves (PRCs). Ideally, we desire a method that has both high precision (few false positives) and high recall (few false negatives), which is represented by a PRC located in the top right region of the plot. Figure 2 shows the precision recall curves for calling true peaks (as defined in Table S2) in the GM12878 10Kb data when the data is down-sampled to different depths. Each panel displays the PRC for peaks called using FitHiC2 as well as HiC-ACT. The shapes indicate where a specific *p*-value threshold for defining FitHiC2/HiC-ACT peaks lies on the curve. For example, with ~0.5 billion raw reads, FitHiC2 achieves a recall of approximately 0.07 and precision near 1 when the significance threshold *p* is between 1.0e-14 and 1.0e-10. However, HiC-ACT with initial *p*-value filtering of 1.0e-3 (blue curve) is able to significantly improve peak classification, achieving recall of approximately 0.36 with negligible loss in precision (0.97) (Figure 2A). As detailed in Table S2, we suggest using a more stringent initial *p*-value filter for data with high sequencing depth (e.g. *p* < 1.0e-6 for data with more than 1 billion raw reads) and using a more lenient initial *p*-value filter for data with shallow sequencing depth (e.g. *p* < 1.0e-3 for data with fewer than 1 billion raw reads). We obtained similar results when the global significance threshold is adjusted (Figures S1-2). HiC-ACT peak calling can be made more conservative by choosing a more stringent initial *p*-value filter. In other words, HiC-ACT allows us to improve precision at the cost of recall (e.g. detection of true peaks), by select a smaller initial *p*-value filter.

**Figure 2.**
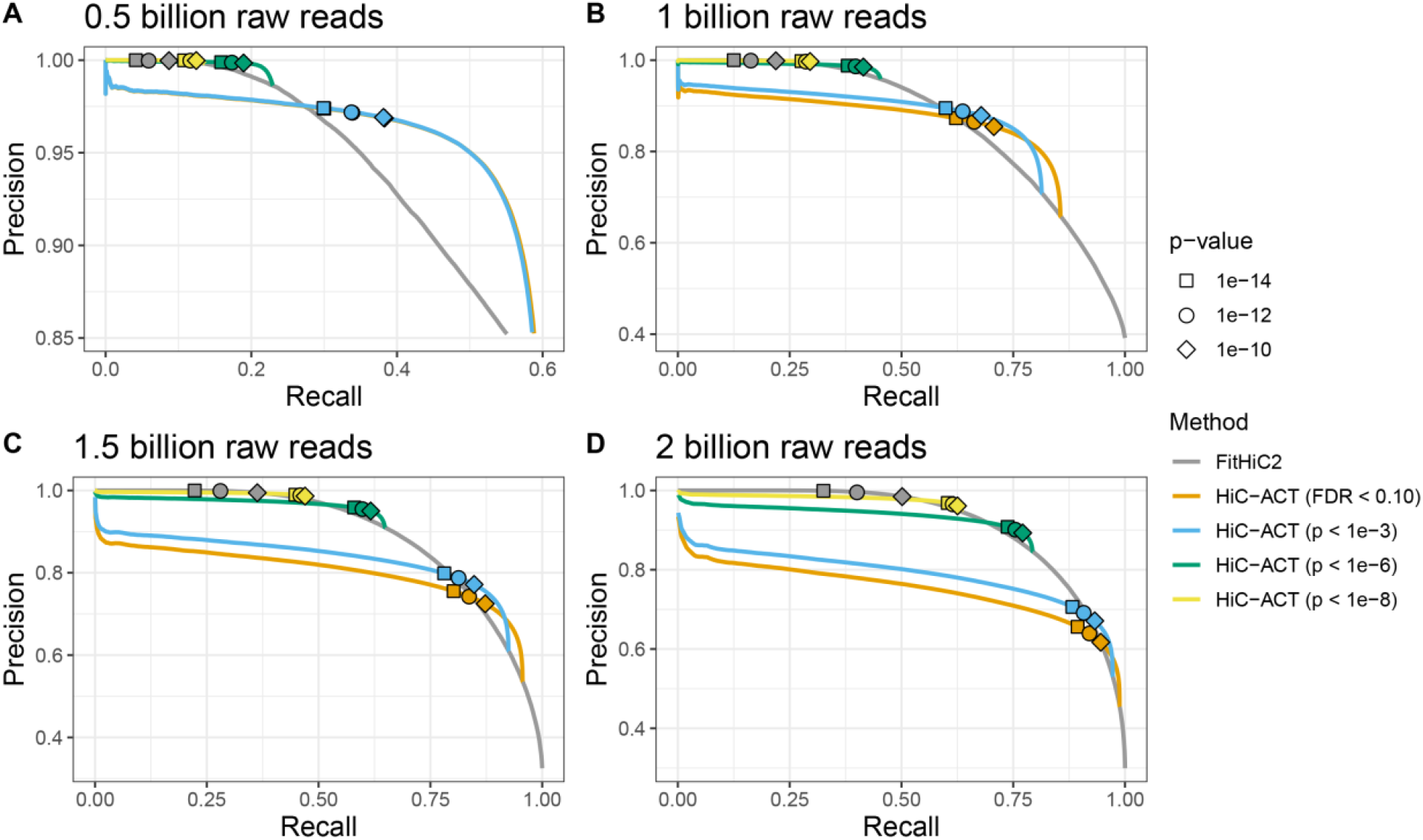
Precision-Recall Curves (PRCs) for calling true peaks. Results for the GM12878 10Kb data down-sampled to ~0.5 billion raw reads (**A**), ~1.0 billion raw reads (**B**), ~1.5 billion raw reads (**C**), and ~2 billion raw reads (**D**) are shown with a global p-value of 1.0e-12 for defining true peaks (based on the full ~4.9 billion raw read data). Each panel displays the PRC for peaks called using FitHiC2 as well as HiC-ACT with various initial filters (FDR/q-value < 0.10 (orange), *p*-value < 1.0e-3 (blue), *p*-value < 1.0e-6 (green), p-value < 1.0e-8 (yellow)). Shapes indicate where a specific *p*-value threshold for defining FitHiC2/HiC-ACT peaks lies on the curve. Note that for ~0.5 billion reads (**A**), the HiC-ACT initial filtering of FDR < 0.10 and *p*-value < 1.0e-3 are nearly identical and thus have overlapping curves.

HiC-ACT also shows improved power to detect significant interactions with low normalized contact frequency. Specifically, we compared the observed to expected contact count ratios between methods for their most significant interactions (ranked *p*-values). Figure 3 shows the distribution of the ratios of the most significant true peaks (significant interactions called in the full data) called by each method in the ~1 billion raw read data. The median ratio for HiC-ACT is ~3.3 (0.5 on the log10 scale) across all top *n* peaks, whereas the median ratio for FitHiC2 decreases from 6 to 4.5 (0.8 to 0.7 on the log10 scale) as the number of top peaks increases. The observed to expected contact count ratios of HiC-ACT are significantly lower than those of FitHiC2 (Wilcoxon test *p* < 2.2e-16) in each case. We reached similar conclusions at other sequencing depths (0.5-2 billion raw reads, data not shown).

**Figure 3.**
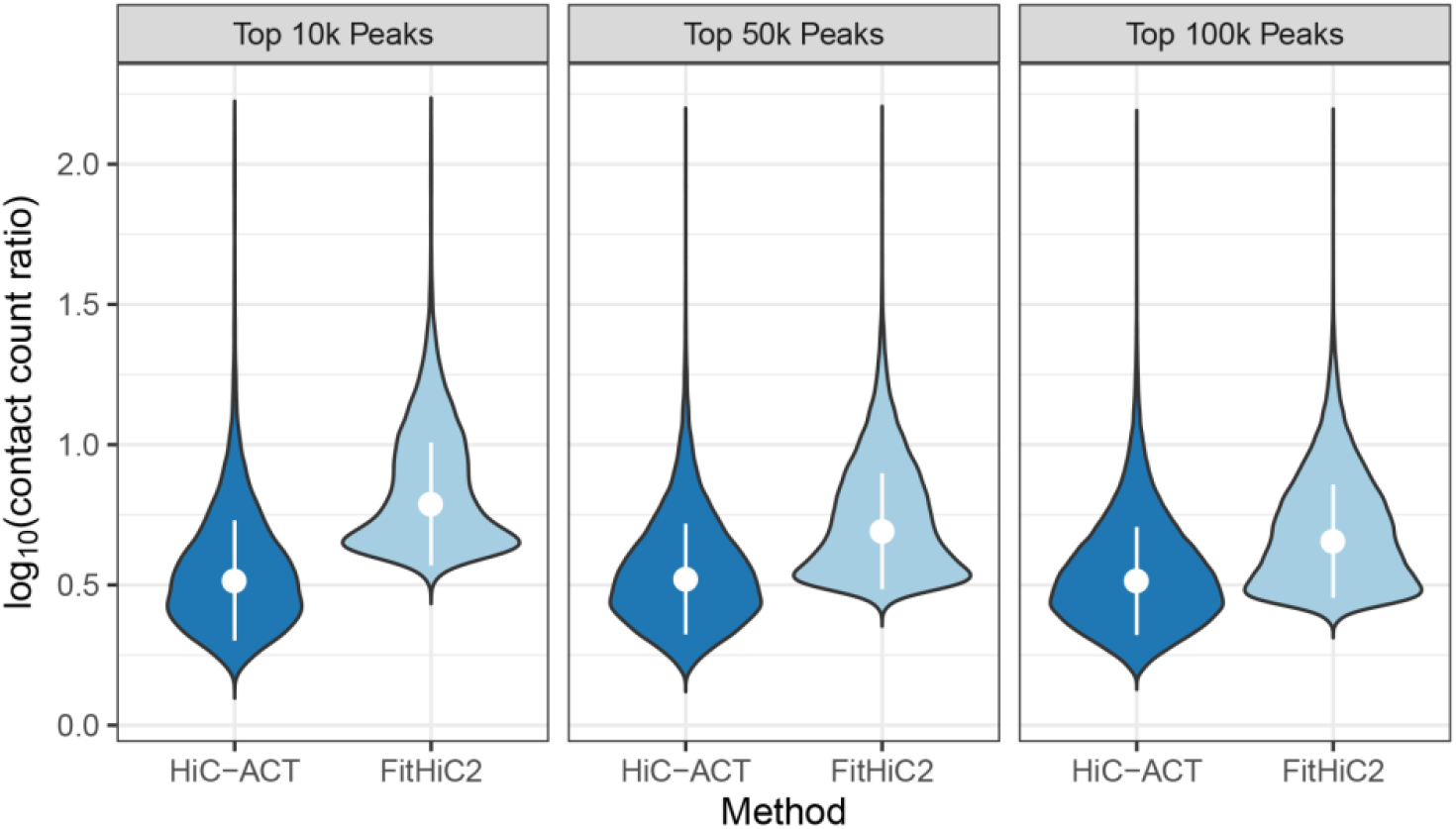
HiC-ACT demonstrates improved power to detect peaks with low normalized counts. Violin plot showing the distribution of the log10 ratio of observed contact counts to expected contact counts for the top true peaks called by HiC-ACT and by FitHiC2 in the GM12878 10Kb data down-sampled to ~1 billion raw reads. Significant interactions were defined using the criteria specified in Table S2, and the true peaks used here further require that the interaction be identified as significant in the full data. Each violin displays the median +/- 1 standard deviation.

### HiC-ACT Identifies Biologically Relevant Interactions

#### GM12878 Roadmap Epigenomics Consortium Enhancers

Using the same GM12878 Hi-C data at 10Kb bin resolution, we compare the peaks called by HiC-ACT and FitHiC2 to typical enhancers (TEs) and super enhancers (SEs) from the Roadmap Epigenomics Consortium^19^. There are 10,335 enhancers in total, 252 of which are super enhancers. First, we identified which peaks have one end overlapping with an enhancer and the other end overlapping with the TSS^20^ of an expressed gene^21^ (FPKM > 1), and defined such peaks as overlapping an enhancer-promoter (E-P) interaction.

At each sequencing depth, we count the total number of super enhancer-promoter (SE-P) interactions (Figure 4A) and typical enhancer-promoter (TE-P) interactions (Figure 4B) identified by each method. HiC-ACT interactions overlap more with SE-P and TE-P interactions than do FitHiC2 interactions. We also count the total number of unique SEs (Figure 4C) and unique TEs (Figure 4D) identified by each method. HiC-ACT is able to capture 90-95% of the SEs and 64-85% of TEs, compared to 77-94% and 38-76%, respectively, captured by FitHiC2. HiC-ACT appears to be less sensitive to sequencing depth than FitHiC2, and shows more significant improvements over FitHiC2 at shallower sequencing depths. Figure 4C-D displays the total counts as well as the odds ratios and corresponding p-values for number of enhancers identified (out of 252 total SEs and 10,335 total TEs).

**Figure 4.**
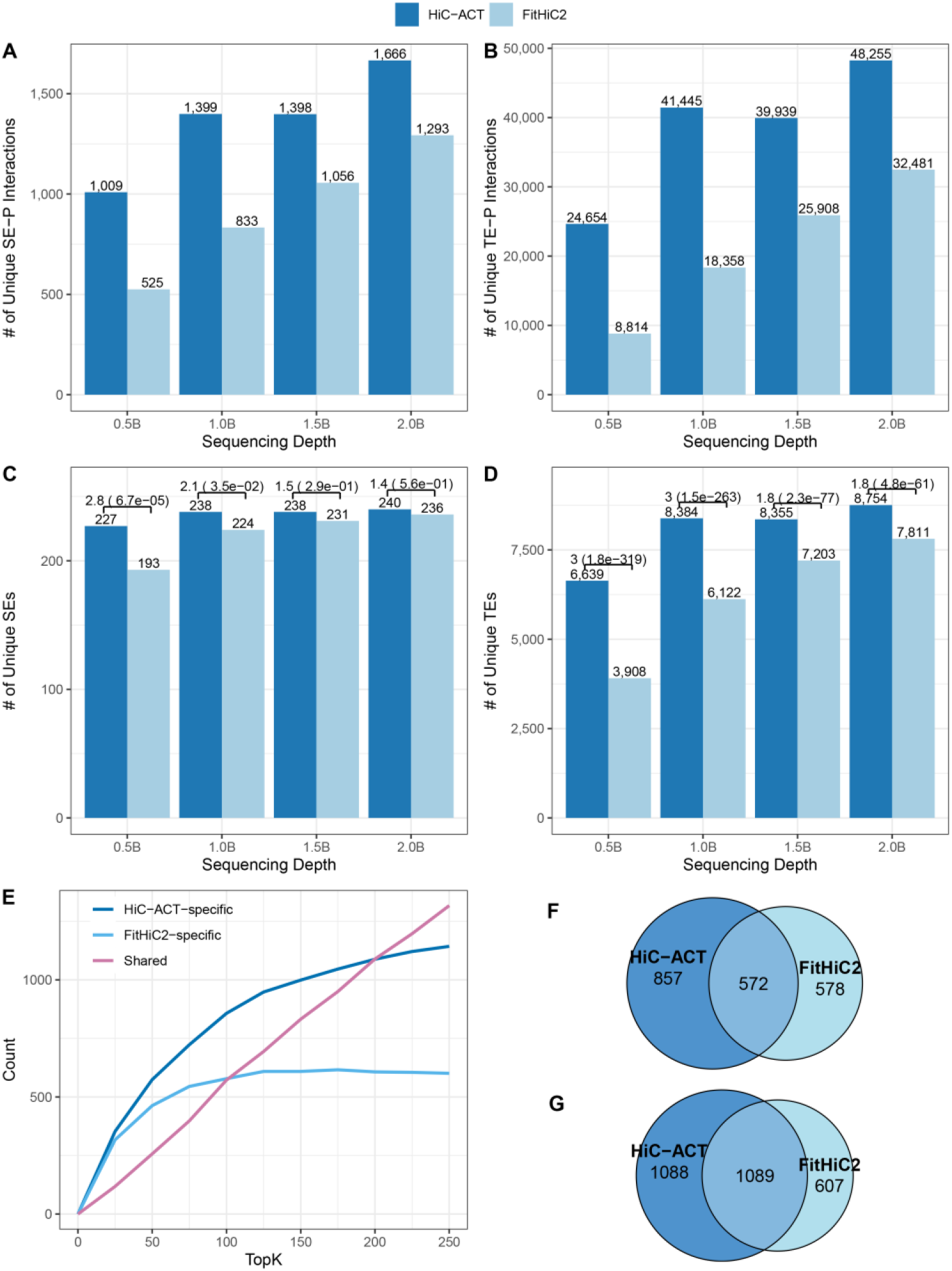
Comparing HiC-ACT and FitHiC2 peak calls with Roadmap Epigenomics Consortium enhancers in GM12878 10Kb Hi-C data. The total number of super enhancer-promoter (SE-P) interactions **(A)** or typical enhancer-promoter (TE-P) interactions **(B)** identified by each method at various sequencing depths (in ~billions of raw reads). The total number of unique super enhancers (SEs) **(C)** or unique typical enhancers (TEs) **(D)** captured is also reported along with the odds ratios and corresponding *p*-values for number of enhancers identified (out of 252 total SEs and 10,335 total TEs). **(E)** The number of HiC-ACT-specific, FitHiC2-specific, and shared interactions overlapping a SE-promoter interaction within a specified number of top peaks (ranked *p*-values) in the GM12878 data down-sampled to ~1 billion raw reads, demonstrating that the most significant interactions identified by each method are different. The breakdown of overlap counts for this example is detailed for the top 100,000 peaks **(F)** and the top 200,000 peaks **(G)**.

Next, we examine the total number of interactions overlapping an E-P interaction within a specified number of top peaks (ranked *p*-values). At all sequencing depths (~0.5-2.0 billion raw reads), we see improved performance of HiC-ACT over FitHiC2 for SE-promoter interactions, and comparable performance between HiC-ACT and FitHiC2 for TE-promoter interactions (Figure S3). Moreover, the most significant interactions identified by each method are different (Figure 4E-G). Figure 4E illustrates the number of HiC-ACT-specific, FitHiC2-specific, and shared peaks that overlap a SE-promoter interaction at various top peaks. For example, out of the top 100,000 peaks called by HiC-ACT and FitHiC2, 2,286 and 1,150 peaks overlap with SE-promoter interactions, respectively. Among them, 857 peaks are HiC-ACT-specific 578 peaks are FitHiC2-specific, and 572 peaks are shared by two methods (Figure 4F). A similar example for the top 200,000 peaks called by each method is displayed in Figure 4G

#### mESC ChIP-seq/ATAC-seq Peaks

We applied HiC-ACT (*h* = 20) to FitHiC2 calls from Hi-C data from mouse embryonic stem cell line (mESC) at 10Kb bin resolution^23^. Significant interactions were defined using the same thresholds as the GM12878 data (observed contact count > 15, expected contact count > 5, the ratio of observed to expected contact counts > 1.5, HiC-ACT filtering *p*-value < 1.0e-6, and global *p*-value < 1.0e-12). By these criteria, HiC-ACT identifies ~1.8 million significant interactions and FitHiC2 identifies ~1 million significant interactions.

We compared these peak calls to mESC ChIP-seq (H3K4me3, H3K4me1, H3K27ac, CTCF) peaks^24–26^ and ATAC-seq peaks^26^. We defined an overlap as a HiC-ACT/FitHiC2 called peak with either 10Kb bin overlapping a ChIP-seq/ATAC-seq peak. Further, we defined a 10Kb bin as an enhancer bin or a promoter bin if it overlaps with a H3K27ac ChIP-seq peak or H3K4me3 ChIP-seq peak and TSS^20^ of an expressed gene^27^ (FPKM > 1), respectively. We defined a HiC-ACT or FitHiC2 identified peak as an enhancer-promoter interaction if one anchor bin is an enhancer bin and the other anchor bin is a promoter bin. We similarly defined enhancer-enhancer and promoter-promoter interactions.

Since HiC-ACT identifies more significant interactions than FitHiC2, we examine the same number of top most significant interactions (ranked *p*-values) called by each method for a fair comparison. The most significant HiC-ACT-specific interactions show higher overlap with enhancer-promoter, enhancer-enhancer, and promoter-promoter interactions than the same number of most significant FitHiC2-specific interactions (Figure 5A). The odds of the most significant HiC-ACT peaks showing overlap with E-P, E-E, or P-P interactions is significantly higher (odds ratio estimate ≈ 1.5, *p* < 7.8e-76) than the odds of the most significant FitHiC2 peaks (Table S4). We see similar results when only considering the 1D overlaps in H3K27ac, H3K4me1, and H3K4me3 ChIP-seq peaks, and a comparable performance between HiC-ACT and FitHiC2 in ATAC-seq peaks and CTCF ChIP-seq peaks (Figure 5B). Table S4 details the number of overlaps displayed by Figure 5 at various numbers of top peaks.

**Figure 5.**
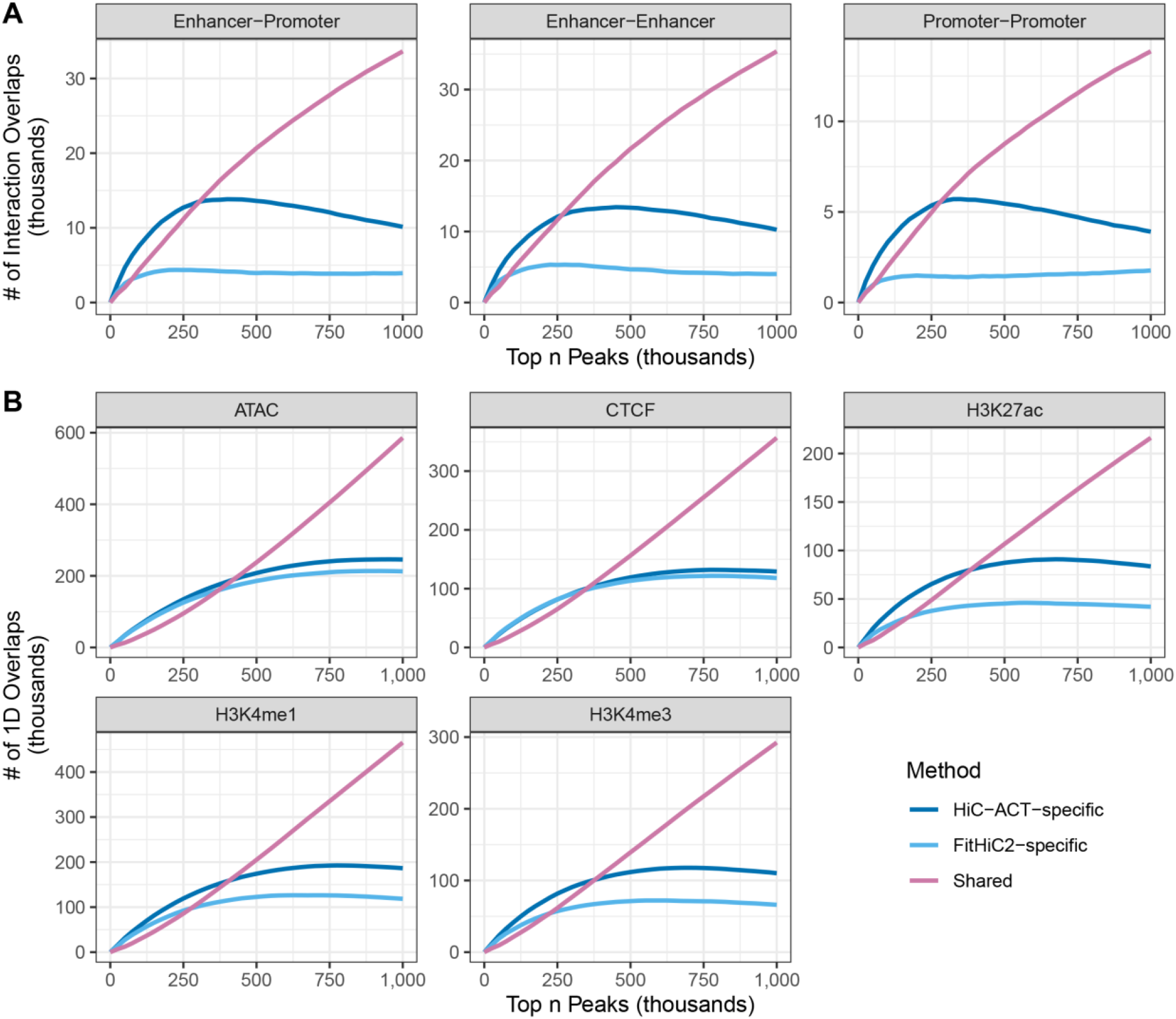
Comparing HiC-ACT and FitHiC peak calls with ChIP-seq and ATAC-seq peaks in mESC data. **(A)** The number of most significant HiC-ACT and FitHiC2 interactions overlapping with an enhancer mark or promoter mark. The most significant HiC-ACT-specific interactions show higher overlap with enhancer-promoter, enhancer-enhancer, and promoterpromoter interactions than the same number of most significant FitHiC2-specific interactions **(B)**. The number of 1D overlaps between a 10Kb bin from most significant HiC-ACT and FitHiC2 interactions and a ChIP-seq/ATAC-seq peak. We see similar results when only considering the 1D overlaps in H3K27ac, H3K4me1, and H3K4me3 ChIP-seq peaks, and a comparable performance between HiC-ACT and FitHiC2 in ATAC-seq peaks and CTCF ChIP-seq peaks. See Table S4 for details relevant to this figure.

#### mESC FANTOM5 and dbSUPER Enhancers

Next, we compared the mESC HiC-ACT and FitHiC2 calls at 10Kb resolution to mESC enhancers cataloged in the FANTOM5^28,29^ database and from the dbSUPER database^30^. FANTOM5 includes 43,662 enhancers and dbSUPER includes 229 super enhancers. For each set of enhancers, we counted how many interactions called by HiC-ACT and FitHiC2 overlap with an enhancer-promoter (E-P) interaction (one end overlapping with an enhancer and the other end overlapping with the TSS^20^ of an expressed gene^27^). The most significant HiC-ACT peaks have approximately 1.4 – 2 times the odds of overlapping an E-P interaction than the same number of most significant FitHiC2 peaks. Figure 6A displays the number of peaks overlapping E-P interactions for each enhancer database and method, as well as the corresponding odds ratio estimates and *p*-values.

**Figure 6.**
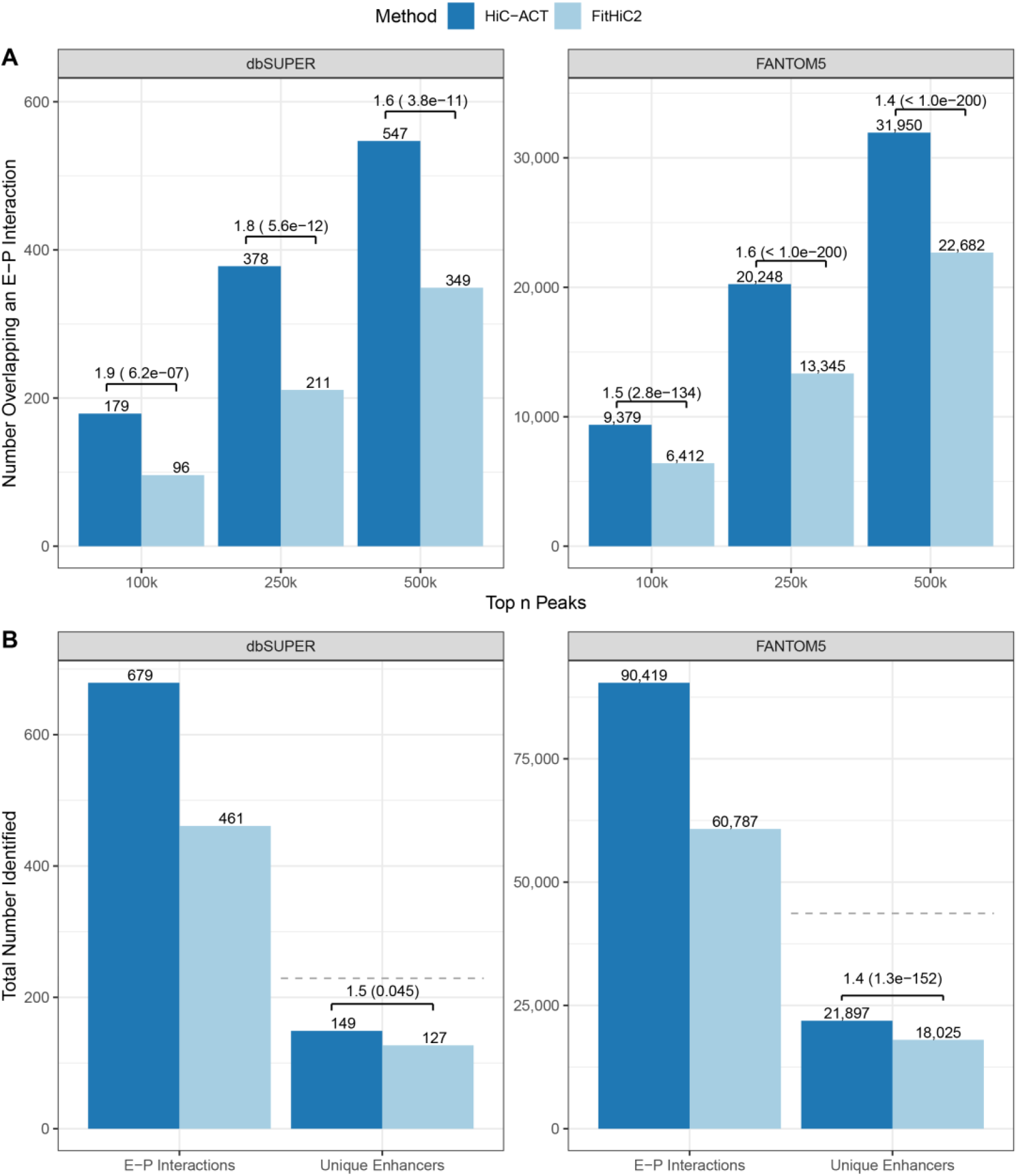
Comparing HiC-ACT and FitHiC peak calls with dbSUPER super enhancers and FANTOM5 enhancers in mESC data. **(A)** The number of interactions called by HiC-ACT and FitHiC2 that overlap with an enhancer-promoter (E-P) interaction (one end overlapping with an enhancer and the other end overlapping with the TSS of an expressed gene). More HiC-ACT peaks overlap with an enhancer than FitHiC2 peaks among their respective most significant 100k, 250k, and 500k interactions. Odds ratios are reported along with their statistical significance (*p*-value). **(B)** The total number of E-P interactions identified by each method for the enhancers in the dbSUPER and FANTOM5 databases. Sine some enhancers may interact with multiple promoters, the total number of unique enhancers among the identified E-P interactions is also reported. The dashed grey line indicates the total number of enhancers in the database.

We next examined the total number of unique enhancer-promoter interactions identified by each method for the enhancers in the dbSUPER and FANTOM5 databases (Figure 6B). HiC-ACT identifies 198 more dbSUPER super enhancer-promoter interactions and 29,632 more FANTOM5 enhancer-promoter interactions than FitHiC2. Further, one enhancer may interact with multiple promoters, so we also report the total number of unique enhancers among the identified E-P interactions. Interestingly, all FitHiC2 identified dbSUPER enhancers and all but 14 FitHiC2 identified FANTOM5 enhancers are also identified by HiC-ACT.

## Discussion

Hi-C has been widely adopted to study chromatin spatial organization with several peak-callers proposed and commonly used to analyze and interpret this data. Here we present HiC-ACT, a method to improve the detection of chromatin interactions by post-processing 3D peak calling results from methods relying on the assumption that pairs of chromatin interactions are statistically independent. HiC-ACT leverages the power of an aggregated Cauchy test to specifically account for the correlation without requiring any information about its structure. We demonstrated that HiC-ACT can improve sensitivity, while maintaining comparable precision. As expected, we observed most pronounced improvement over FitHiC2 when sequencing depth is less than one billion reads, which is the typical depth for the vast majority of Hi-C data generated to date. Even with increasing sequencing depth anticipated in some future Hi-C studies, we consider HiC-ACT useful as it will allow more powerful 3D peak calling at finer resolution (e.g., 5Kb or even 1Kb resolution, particularly when cut with the appropriate restriction enzymes such as the 4 base pair cutter MboI or DpnII).

Although HiC-ACT can theoretically be applied to HiCCUPS output, we consider such application inappropriate due to the intrinsic nature of HiCCUPS to call summits in peak regions. HiCCUPS contrasts each chromatin interaction pair with its local neighborhood; however, our goal is to call peaks by borrowing information from the neighborhood.

HiC-ACT is computationally efficient and scalable. HiC-ACT can process 25,000 pairwise interactions in ~6 minutes with ~2GB memory and 0.5 million pairwise interactions in ~2 hours with ~30GB memory, using a 2.50 and 3.40 GHz Intel processor, respectively. Note that chromosome 1 has ~90,000 and ~168,000 pairwise interactions, at 10Kb resolution, passing the suggested initial *p*-value filter in the ~0.5 and ~1 billion raw reads GM12878 Hi-C data, respectively.

Future work may involve fine tuning the smoothing parameter, particularly for 1Kb bin resolution Hi-C data, and investigating different weight functions.

By identifying statistically significant long-range interactions with enhanced statistical power and improved computationally efficiency, HiC-ACT can improve our knowledge regarding regions with regulatory potential, and aid to establish links between regulatory regions and their target genes. We anticipate HiC-ACT will become a convenient tool for many researchers.

## Appendix

## Claim

When only one *p*-value is considered 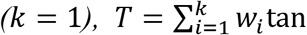 follows a Cauchy distribution (*x*_0_ = 0, *γ* = *w*_1_) under the null hypothesis that *p*_1_ is uniformly distributed between 0 and 1.

*Proof:*

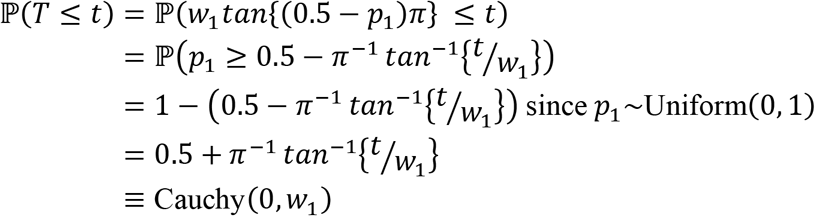

## Description of Supplemental Data

Supplemental Data include three figures and four tables, one of which is included as an Excel file.

## Acknowledgements

This material is based upon work supported by the National Science Foundation Graduate Research Fellowship Program under Grant No. DGE-1650116. TL and YL are partially funded by R01 HL129132 (awarded to YL). YL is additionally supported by R01 GM105785 and P50 HD103573. AA and MH are partially funded by NIH grants U54DK107977 and UM1HG011585 (awarded to MH).

## Web Resources

HiC-ACT software can be freely downloaded at https://github.com/tmlagler/hicACT or through our lab website at https://yunliweb.its.unc.edu/hicACT/.

## Data and Code Availability

The authors declare no competing interests.

## Accession Numbers

This paper did not generate any data sets.

