## Supplemental Figures/Tables for "HiC-ACT: Improved Detection of Chromatin Interactions from Hi-C Data via Aggregated Cauchy Test"

### Supplemental Data

**Figure S1. Precision-Recall Curves for calling true peaks.** Results for the GM12878 10Kb data[3] down-sampled to ~0.5 billion raw reads (**A**), ~1.0 billion raw reads (**B**), ~1.5 billion raw reads (**C**), and ~2 billion raw reads (**D**) are shown with a global p-value of 1.0e-10 for defining true peaks (based on the full ~4.9 billion raw read data). Each panel displays the precision recall curve for peaks called using FitHiC2 as well as HiC-ACT with various initial filters (FDR/q-value < 0.10 (orange), *p*-value < 1.0e-3 (blue), *p*-value < 1.0e-6 (green), *p*-value < 1.0e-8 (yellow)). Shapes indicate where a specific p-value threshold for defining FitHiC2/HiC-ACT peaks lies on the curve. Note that for ~0.5 billion reads (**A**), the HiC-ACT initial filtering of FDR < 0.10 and *p*-value < 1.0e-3 are nearly identical and thus have overlapping curves.

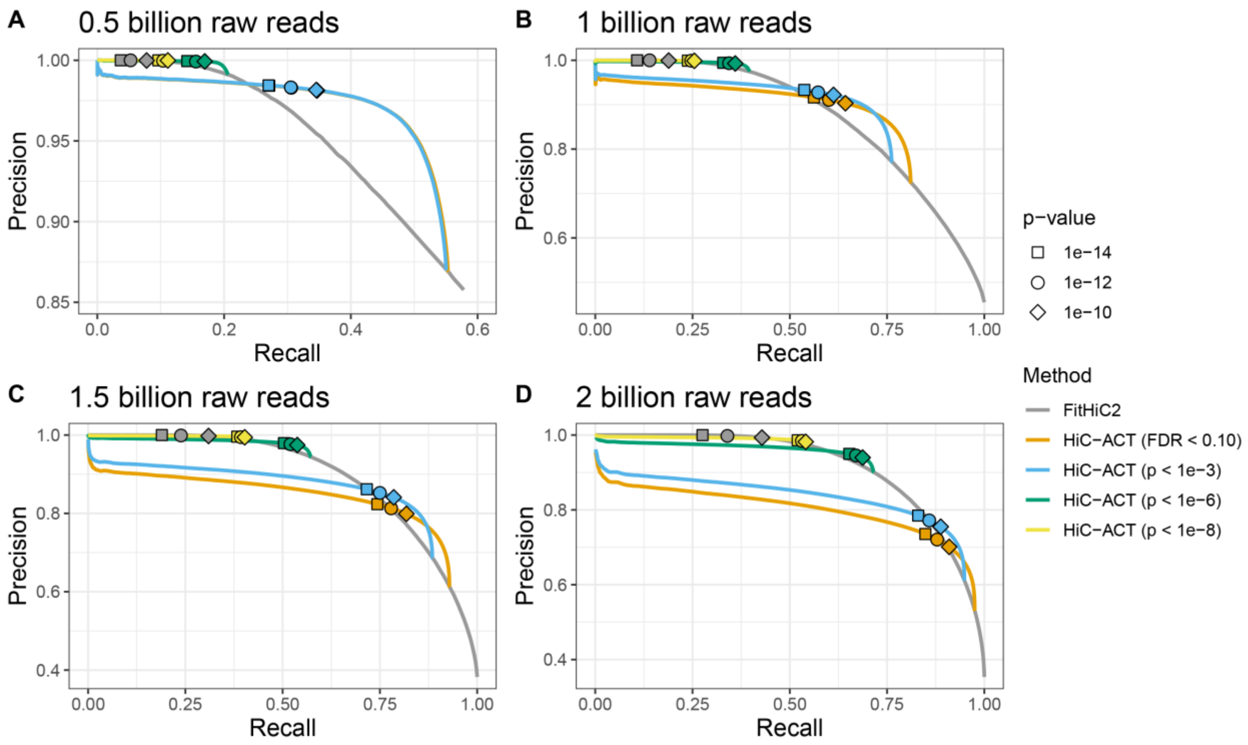

**Figure S2. Precision-Recall Curves for calling true peaks.** Results for the GM12878 10Kb data[3] down-sampled to ~0.5 billion raw reads (**A**), ~1.0 billion raw reads (**B**), ~1.5 billion raw reads (**C**), and ~2 billion raw reads (**D**) are shown with a global p-value of 1.0e-14 for defining true peaks (based on the full ~4.9 billion raw read data). Each panel displays the precision recall curve for peaks called using FitHiC2 as well as HiC-ACT with various initial filters (FDR/q-value < 0.10 (orange), *p*-value < 1.0e-3 (blue), *p*-value < 1.0e-6 (green), *p*-value < 1.0e-8 (yellow)). Shapes indicate where a specific p-value threshold for defining FitHiC2/HiC-ACT peaks lies on the curve. Note that for ~0.5 billion reads (**A**), the HiC-ACT initial filtering of FDR < 0.10 and *p*-value < 1.0e-3 are nearly identical and thus have overlapping curves.

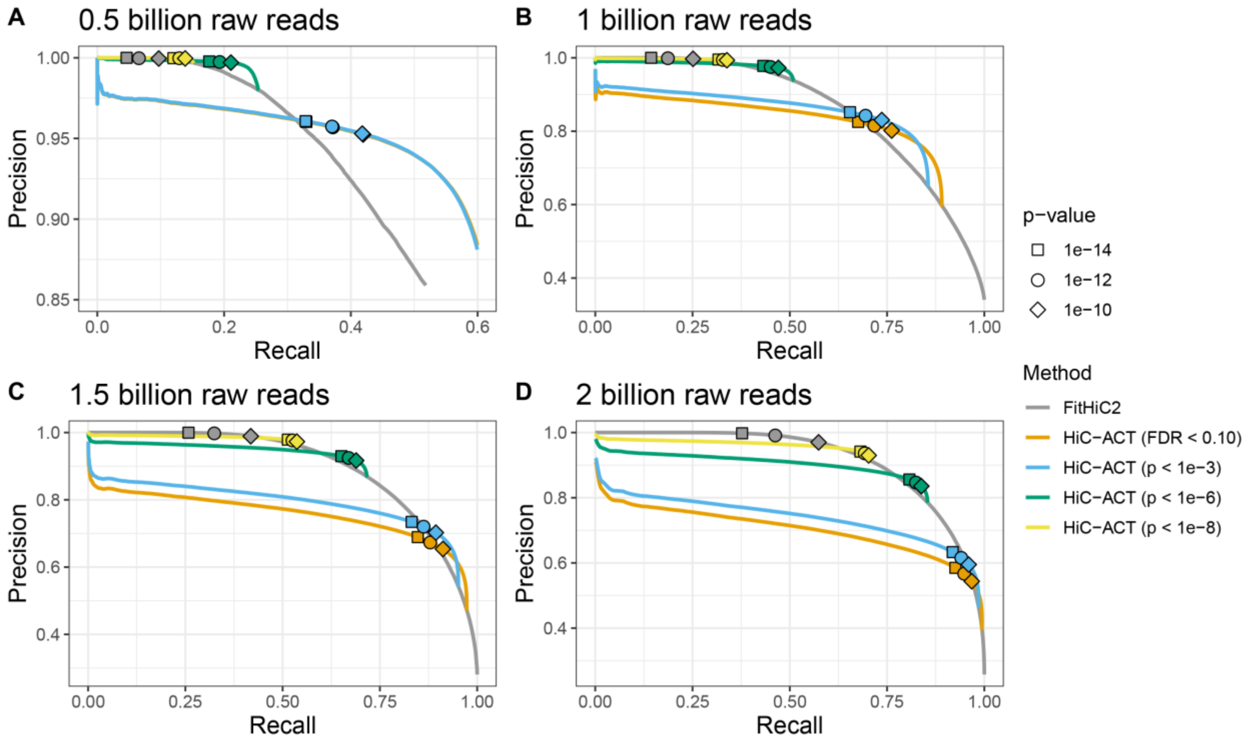

**Figure S3. Comparing HiC-ACT and FitHiC2 peak calls with Roadmap Epigenomics Consortium[4] enhancers in GM12878 10Kb Hi-C data.** HiC-ACT-specific, FitHiC2-specific, and shared interactions overlapping a super enhancer (SE)-promoter interaction (left column) or a typical enhancer (TE)-promoter interaction (right column) within a specified number of top peaks (ranked *p*-values) in the GM12878 data down-sampled to ~0.5-2 billion raw reads (rows). The most significant HiC-ACT-specific interactions show higher overlap with SE-promoter interactions than the same number of most significant FitHiC2-specific interactions. Performance between the two methods is comparable for overlap with TE-promoter interactions. In each case, we observe that the most significant interactions identified by each method are different.

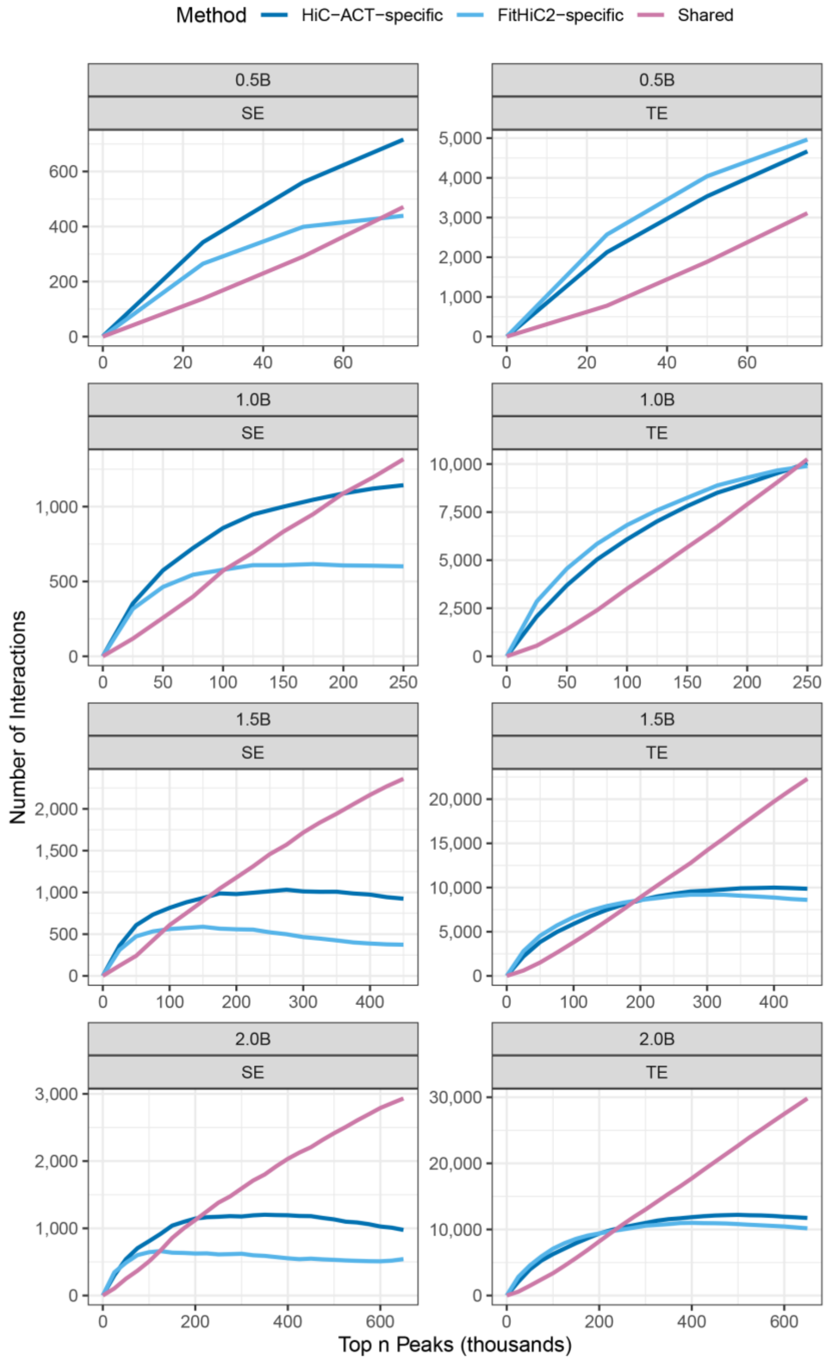

**Table S1. Recommendations for selecting a HiC-ACT smoothing parameter (*h*) following the HiCRep[1] method based on data resolution.**

| Data Resolution (Kb) | Smoothing Parameter (*h*) | From HiCRep |
|---|---|---|
| 5 | 40 | no |
| 10 | 20 | yes |
| 20 | 12 | no |
| 25 | 11 | yes |
| 40 | 5 | yes |

**Table S2. Recommendations for defining HiC-ACT peaks at various sequencing depths (in approximate billions of raw reads).** minCC is the minimum number of observed contact counts and minEC is the minimum number of expected contact counts. Peaks are also required to have a ratio of observed to expected contact counts of at least 1.5. Initial *p* filtering refers to which *p*-values are to be considered for smoothing. The global *p* threshold determines if the smoothed *p*-value is significant. The number of significant FitHiC2 contacts depends on sequencing depth of Hi-C data[2]. Consequently, we allowed the global *p* thresholds to become more lenient as the sequencing depth decreased. The results are robust to minor adjustments in the global *p* threshold, as shown in the precision recall curves (Figure 2, Figures S1-2).

| Sequencing Depth (unit: billion) | minCC | minEC | Initial *p* filtering | Global *p* threshold |
|---|---|---|---|---|
| 0.5 | 5 | .5 | 1.0e-3 | 1.0e-11 |
| 1 | 5 | 1 | 1.0e-3 | 5.0e-12 |
| 1.5 | 5 | 1.5 | 1.0e-6 | 3.3e-12 |
| 2 | 5 | 2 | 1.0e-6 | 2.5e-12 |
| 4.9 | 15 | 5 | *NA* | 1.0e-12 |

**Table S4. Comparing HiC-ACT and FitHiC peak calls with ChIP-seq and ATAC-seq peaks[5–7] in mESC data.** The number of the 100k **(A)**, 250k **(B),** 500k **(C)**, and 750k **(D)** most significant HiC-ACT and FitHiC2 interactions which have each end overlapping with an enhancer mark or promoter mark followed by the number of 1D overlaps between a 10Kb bin from most significant HiC-ACT and FitHiC2 interactions and a ChIP-seq/ATAC-seq peak. The most significant HiC-ACT-specific interactions show higher overlap with enhancer-promoter (E-P), enhancer-enhancer (E-E), and promoter-promoter (P-P) interactions than the same number of most significant FitHiC2-specific interactions. Odds ratio estimates and corresponding *p*-values for the proportion of total HiC-ACT (HiC-ACT-specific + Shared) and total FitHiC2 (FitHiC2-specific + Shared) most significant interactions overlapping an E-P, E-E, or P-P interaction out of the total number of top peaks examined are reported. The odds of the most significant HiC-ACT peaks showing overlap with E-P, E-E, or P-P interactions is significantly greater than the odds of the most significant FitHiC2 peaks. We see similar results when only considering the 1D overlaps in H3K27ac, H3K4me1, and H3K4me3 ChIP-seq peaks, and a comparable performance between HiC-ACT and FitHiC2 in ATAC-seq peaks and CTCF ChIP-seq peaks.

**A. Top 100k Peaks**

| | HiC-ACT-specific | FitHiC2-specific | Shared | OR Estimate | p-value |
|---|---|---|---|---|---|
| Enhancer-Promoter | 7,653 | 3,508 | 4,446 | 1.59 | 2.4e-210 |
| Enhancer-Enhancer | 7,366 | 4,123 | 4,808 | 1.41 | 1.6e-123 |
| Promoter-Promoter | 3,321 | 1,304 | 1,991 | 1.65 | 2.8e-110 |
| CTCF | 40,500 | 41,497 | 22,221 | - | - |
| H3K4me1 | 58,090 | 47,918 | 28,458 | - | - |
| H3K4me3 | 41,673 | 31,937 | 21,446 | - | - |
| H3K27ac | 34,462 | 22,261 | 16,935 | - | - |
| ATAC | 63,115 | 60,400 | 30,722 | - | - |

**B. Top 250k Peaks**

| | HiC-ACT-specific | FitHiC2-specific | Shared | OR Estimate | p-value |
|---|---|---|---|---|---|
| Enhancer-Promoter | 12,723 | 4,351 | 11,174 | 1.6 | < 3.8e-271 |
| Enhancer-Enhancer | 12,077 | 5,293 | 11,632 | 1.44 | 3.8e-271 |
| Promoter-Promoter | 5,354 | 1,457 | 4,961 | 1.63 | 6.3e-208 |
| CTCF | 81,709 | 82,037 | 65,236 | - | - |
| H3K4me1 | 119,278 | 92,304 | 85,667 | - | - |
| H3K4me3 | 81,703 | 57,354 | 61,606 | - | - |
| H3K27ac | 65,344 | 37,676 | 48,736 | - | - |
| ATAC | 134,529 | 125,494 | 94,920 | - | - |

### Table S4 – Continued

#### C. Top 500k Peaks

| | HiC-ACT-specific | FitHiC2-specific | Shared | OR Estimate | p-value |
|---|---|---|---|---|---|
| Enhancer-Promoter | 13,603 | 3,943 | 20,684 | 1.42 | $<$ 2.4e-288 |
| Enhancer-Enhancer | 13,357 | 4,662 | 21,661 | 1.36 | 2.4e-288 |
| Promoter-Promoter | 5,454 | 1,449 | 8,755 | 1.4 | 4.7e-149 |
| CTCF | 119,254 | 113,388 | 156,336 | - | - |
| H3K4me1 | 174,257 | 122,681 | 205,120 | - | - |
| H3K4me3 | 111,723 | 71,162 | 139,261 | - | - |
| H3K27ac | 87,284 | 45,221 | 106,722 | - | - |
| ATAC | 208,023 | 185,826 | 237,686 | - | - |

#### D. Top 750k Peaks

| | HiC-ACT-specific | FitHiC2-specific | Shared | OR Estimate | p-value |
|---|---|---|---|---|---|
| Enhancer-Promoter | 12,071 | 3,860 | 27,776 | 1.27 | 8.9e-218 |
| Enhancer-Enhancer | 12,120 | 4,177 | 29,222 | 1.25 | 1.6e-195 |
| Promoter-Promoter | 4,711 | 1,584 | 11,555 | 1.24 | 7.8e-76 |
| CTCF | 131,698 | 121,595 | 255,400 | - | - |
| H3K4me1 | 192,246 | 126,146 | 335,221 | - | - |
| H3K4me3 | 117,364 | 70,952 | 217,335 | - | - |
| H3K27ac | 90,146 | 44,743 | 163,132 | - | - |
| ATAC | 240,532 | 210,032 | 404,924 | - | - |
